# In vivo evidence for bleb-induced survival signaling in metastatic melanoma

**DOI:** 10.64898/2026.01.05.697601

**Authors:** Andrew D. Weems, Md Torikul Islam, Hazel Borges, Clarence Yapp, Lizbeth Perez-Castro, Pedro Nogueira, Jinlong Lin, Zoltan Maliga, Alecxander J. Lewis, Camille Kutter, Mark Mannino, Tuulia Vallius, Sandro Santagata, Peter Sorger, Maralice Conacci-Sorrell, Kevin M. Dean, Gaudenz Danuser

**Author notes:** Correspondence to Andrew Weems.

## Abstract

Bleb signaling is a cellular process in which pressure-driven plasma membrane protrusions generate localized micron-scale membrane curvature that recruits cytosolic septin complexes, assembling signaling hubs that promote cell survival^1^. Our prior work showed that this morphology-encoded signaling pathway is necessary to sustain anchorage-independent survival of BRAF and NRAS mutant melanoma cells *in vitro*. However, whether bleb-induced signaling occurs *in vivo* and contributes to cancer progression is unclear. Here, we develop complementary *in vivo* and *ex vivo* assays spanning human patient samples and mouse xenograft models to mechanistically interrogate bleb signaling directly in physiologic contexts. We observe that bleb-associated septin hubs are exclusively formed in poorly adherent amoeboid tumor cells at the invasive margin, within malignant effusions, and at distant metastatic sites, while well-adhered cells in the tumor interior show no signs of septin hub formation. Accordingly, pharmacological pathway disruption specifically kills disseminated tumor cells within these low-adhesion microenvironments while having no appreciable effect on adhered cells in the tumor interior, resulting in reduced metastatic burden and delayed disease recurrence *in vivo*. This work confirms septin-mediated bleb signaling as a previously unrecognized vulnerability of disseminated cancer cells and demonstrates that this morphology-encoded pathway operates *in vivo* to support disease progression, preserving cancer cell viability under conditions in which cell survival signals from the environment are muted. These findings suggest novel opportunities to target survival signaling in micrometastatic disseminated cancer cells.

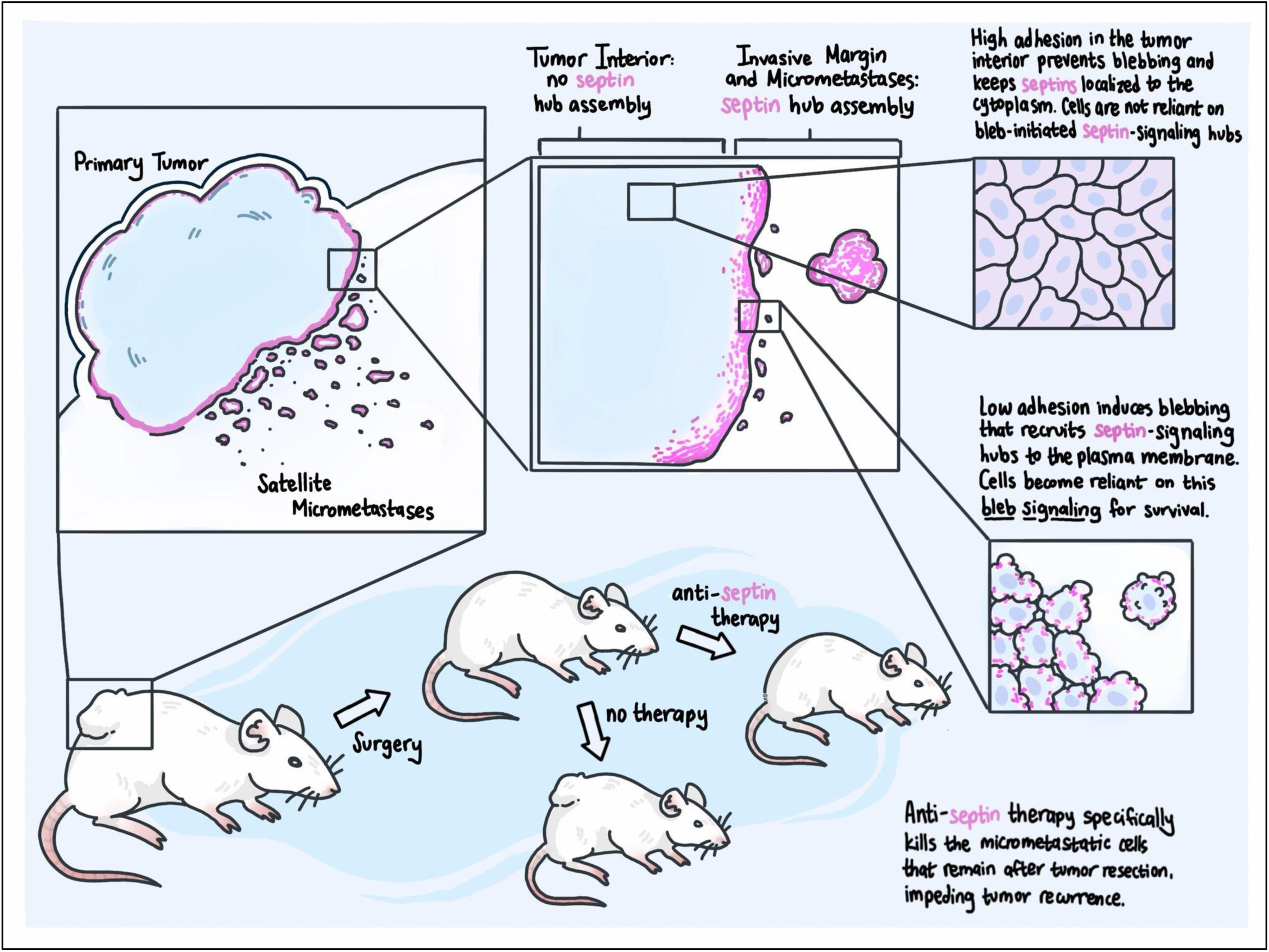

## MAIN

Pronounced shifts in cellular morphology have long been associated with malignancy, with the amoeboid cancer cell morphotype being of great and increasing interest due to its association with aggressive metastatic behavior^2^. The surfaces of these amoeboid cancer cells are densely decorated with blebs – round dynamic protrusions of the plasma membrane that are established by intracellular pressure in locations of small defects in membrane-cortex attachment^3,4^. Though blebs are widely recognized as key drivers of amoeboid cell motility, their potential role in enabling additional functional transformations that confer aggressive cancer cell behavior has remained poorly defined^5–7^. Our recent work has added the formation of potent signaling hubs to the repertoire of established bleb functions.^1,8,9^. In this process bleb-generated plasma membrane curvature activates the assembly of curvature-sensing septin scaffolds^10^ that organize and upregulate plasma membrane bound signaling factors. In cultured melanoma cells we find that these hubs concentrate mutant NRAS and upregulate signal transduction to downstream MAPK and PI3K/AKT pathways. We have shown that blebs and the septin structures they assemble are a requirement for NRAS(Q61R) to assume its oncogenic function as a driver of anchorage-independent cell survival – a hallmark of cancers with high metastatic propensity.

The specifics of this mechanism are compelling, as the pathway seems to specifically rely on the activity of the septin cytoskeleton, suggesting a constrained molecular dependency that may represent a content-specific therapeutic vulnerability. Moreover, the wide variety of cancers types that display amoeboid morphotypes raises the possibility that a therapeutic approach targeting this vulnerability might be applicable beyond melanoma^2^. The results underpinning these ideas, however, have up until now been limited to *in vitro* experiments performed on immortalized cancer cell lines either in suspension or embedded in biomimetic 3D hydrogels^1^. To test the relevance of these findings in the context of *in vivo* disease progression, we performed a series of new experiments that rely on assays with both human clinical samples and murine xenograft models of melanoma, testing the roles bleb signaling plays in various stages of formation of metastatic cancer.

### Melanoma cells activate bleb signaling *in vivo*

Cancer cells *in vitro* specifically rely on bleb signaling to generate survival signals when they reside in environments lacking opportunities for stable attachment^1^. To test whether this morphology-encoded survival pathway also operates in vivo, we sought human tumor contexts in which cancer cells experience sustained detachment. While several “low attachment” environments exist in the body (including blood and lymph), isolated detached cancer cells are generally rare, making the harvesting of experimentally meaningful samples difficult. A notable exception is provided by malignant effusions, from which cytology blocks are routinely prepared following fine needle aspiration for clinical diagnosis^11,12^. These abnormal fluid-filled cavities, which often develop in patients with advanced metastatic disease, accumulate large numbers of suspended cancer cells (present individually and in small clusters) and therefore provide a physiologically relevant and experimentally tractable source of non-adherent cells.

We obtained cytology blocks from patients with advanced melanoma collected from three different settings: cutaneous melanoma metastatic to the salivary gland, mucosal melanoma present in an abdominal (peritoneal) effusion, and vulvar melanoma present in a lung (pleural) effusion. Immunohistochemistry (IHC) staining for SEPT2, a marker we previously validated as labeling bleb-generated septin signaling hubs^1^, revealed septin-containing scaffolds enriched at the plasma membrane of rounded, bleb-rich cancer cells (Fig 1A-C). To examine these structures at higher resolution, we selected the specimen that most clearly maintained a blebby PM morphology post-fixation. This HER2-amplified mucosal melanoma sample was de-waxed, immunofluorescently stained against both SEPT2 and HER2 (to confirm cancer identity), and imaged by laser scanning confocal microscopy. The resulting micrographs confirmed the conclusion from IHC that HER2-positive cancer cells enrich septin localization in the blebby regions of the PM (Fig 1D). Interestingly, septin signals closely colocalized with HER2 (Fig S1 and Movie 1), consistent with prior observations that septin scaffolds organize HER2 at the plasma membrane in adherent epithelial cancers^13^ and extending this phenomenon to detached amoeboid melanoma cells via bleb-associated septin hub assembly. Together, these cytology-based observations indicate that the formation of bleb signaling hubs is not restricted to detached melanoma cells *in vitro*, but also occurs in human cancers *in vivo* under conditions of sustained loss of adhesion.

**Figure 1:**
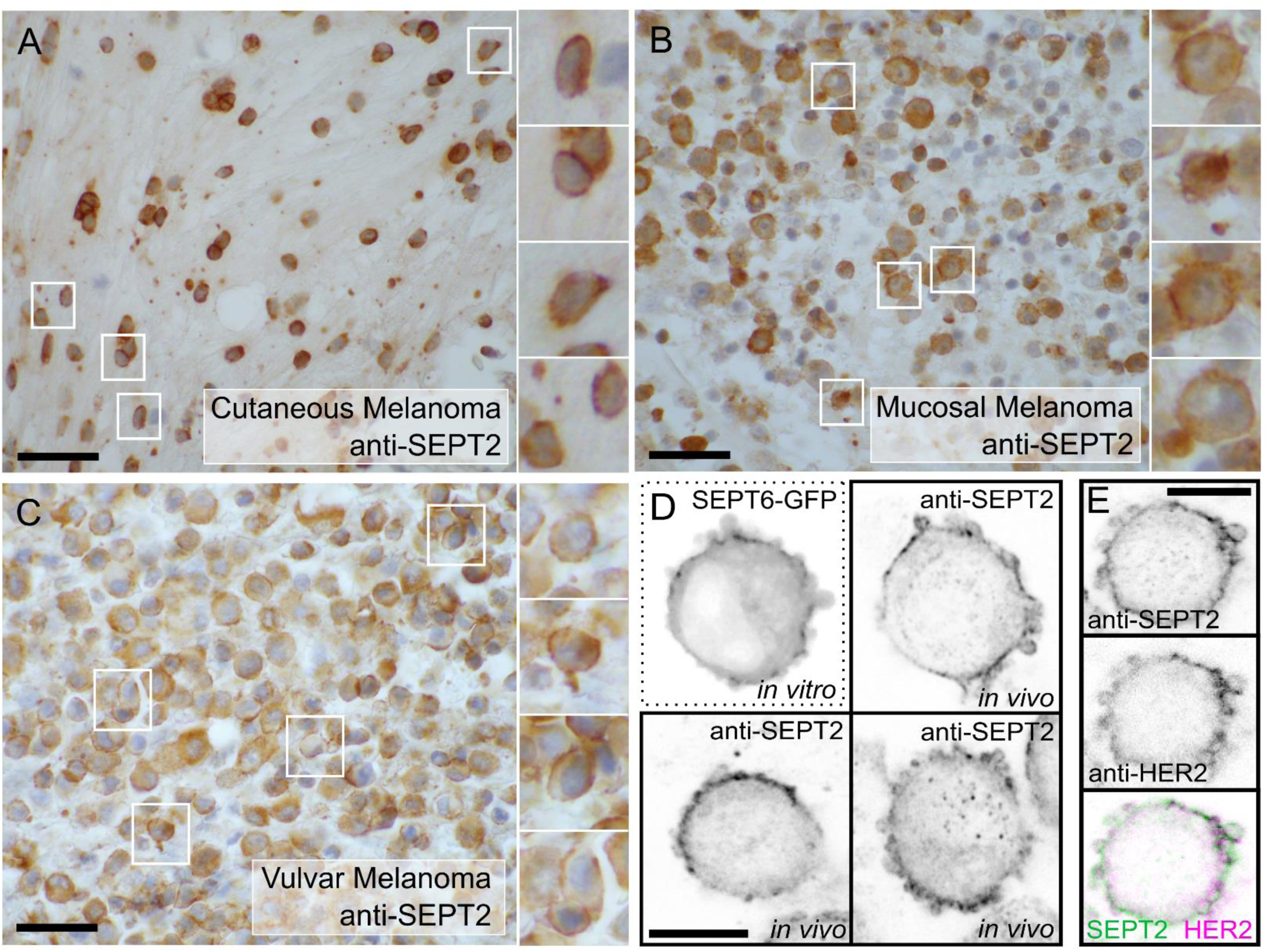
Detached metastatic melanoma cells activate bleb signaling in malignant pleural effusions. **a-c**) Cell block preparations from three different melanoma patients suffering from malignant effusions of the salivary gland (**a**), peritoneum (**b**), and pleura (**c**), immunostained for SEPT2. All show solitary melanoma cells with strong localization of SEPT2 at the plasma membrane. Insets to the right magnify the regions marked with white boxes. **d)** Cells from the mucosal melanoma peritoneal effusion sample (**b**) were immunofluorescently stained for SEPT2 and imaged using Laser Scanning Confocal Microscopy (LSCM) to show finer features of the plasma membrane and septin localization. An *in vitro* MV3 melanoma cell expressing SEPT6-GFP and imaged while suspended in soft collagen is shown in the upper-left panel marked by the dotted line for comparison. Scalebars for indicate 50 μm for **a-c**, and 20 μm for **d**.

### Melanoma cells specifically activate bleb signaling within the invasive margin and proximal stroma of human primary tumors

With cytology specimens supporting the occurrence of bleb signaling in metastatic cells within malignant effusions, we next sought to determine whether bleb signaling hubs could be observed in other clinically-relevant microenvironmental contexts. Bleb signaling might be especially important at the invasive margin and in the proximal stroma, microenvironments in which nascently metastatic cancer cells migrating away from the primary tumor begin to experience loss of attachment and thus may increasingly depend on cell autonomous survival cues^2,14,15^. Previous research has shown that this transition in attachment strength tends to shift cells away from mesenchymal morphologies towards the more rounded and amoeboid morphotype^2^, further indicating that bleb signaling might play a critical role in these challenging conditions of metastasis initiation.

To test this, we secured several primary tumor biopsies from patients suffering from a variety of skin cancers, including nodular acral melanoma, mucosal melanoma, superficial cutaneous melanoma, and Merkel cell carcinoma. In all cases we ensured that samples included not only intratumoral regions, but also invasive margins and surrounding stroma, allowing for comparisons of septin localization in these different areas. After staining samples using the SEPT2 antibody described above, a stark contrast emerged between septin localization within cancer cells found in the inner tumor vs. the invasive margin. The ostensibly poorly-attached cells at the invasive margin tended to display amoeboid morphologies with strong septin enrichment at the PM, while cells in the high-attachment tumor interior displayed more diffuse septin signals throughout the cell with little-to-no enrichment at cell edges (Fig 2A-D). This aligned with our previously published *in vitro* results showing that melanoma cell detachment initiates blebbing that recruits membrane curvature sensing septins to the PM^1^. Given that this septin accumulation results in the activation of survival signaling *in vitro*, we hypothesized that the observed transition to blebby morphotypes with septin enriched PMs may similarly promote the survival of these invasive cells *in vivo*.

**Figure 2:**
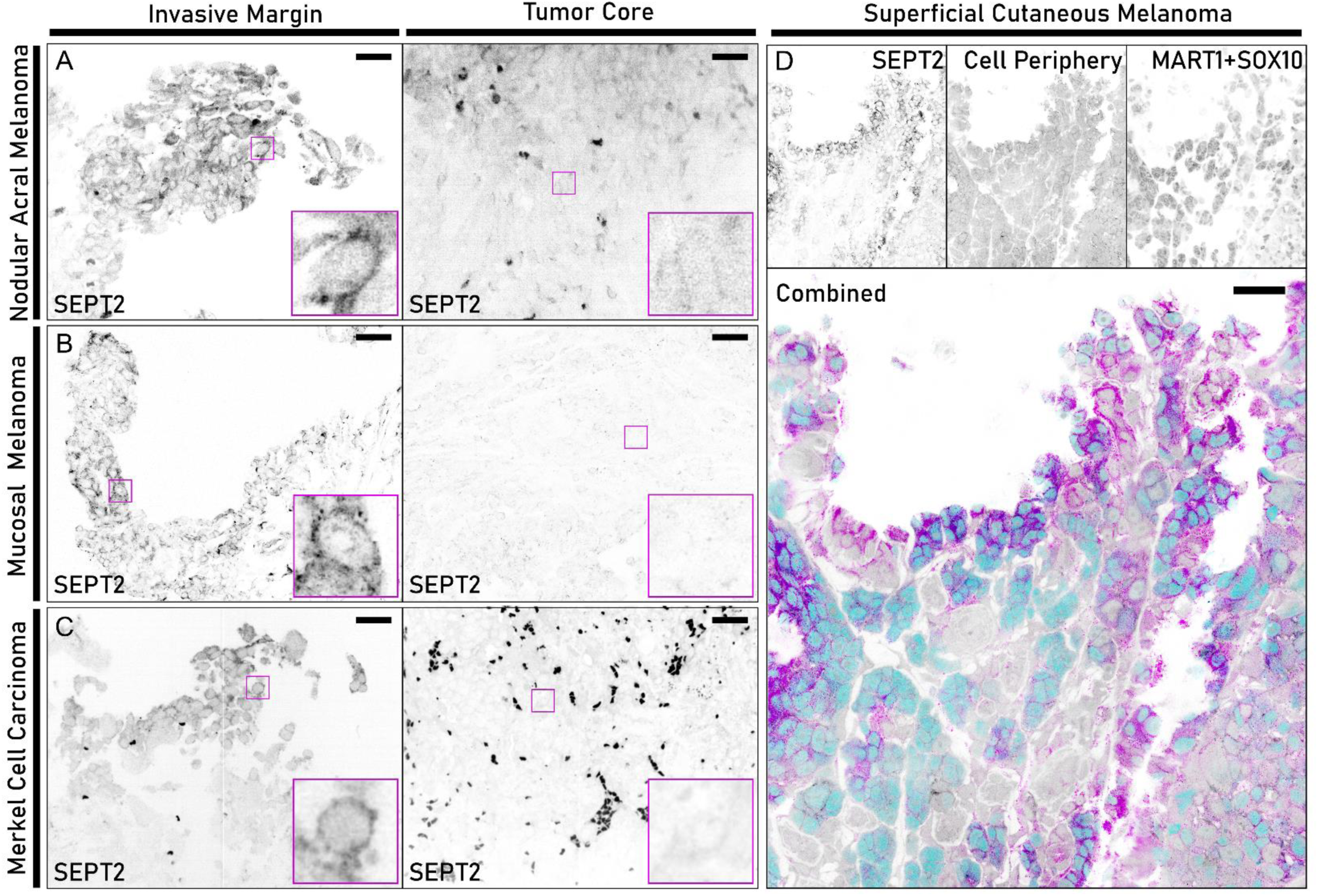
Melanoma cells specifically activate bleb signaling at the invasive margin. Human primary tumor samples from different skin cancers showing regional trends in the activation of bleb signaling via septin localization. **(a-c)** Comparisons of bleb signaling activation between cells in the invasive margin (left) and tumor interior (right) of 3 different skin cancer varieties with magnified insets at the lower left. Poorly-adhered cells at the invasive margin bear rounded blebby morphologies and PM-localized septins (shown with immunofluorescent staining of SEPT2) indicative of bleb signaling, while well-adhered cells in the nearby tumor interior do not. **(d)** Large field of view from a superficial cutaneous melanoma showing activation of bleb signaling specifically at the edge of the tumor. Cells immunofluorescently stained for SEPT2 to show septin localization (magenta), a combination of MART1 and SOX10 to indicate melanocyte lineage (cyan), and a combination of MHC-I, MHC-II, E-cadherin, and β-catenin to mark the periphery and cytoplasm of all cells (gray). **a** and **c** were cleared intact specimens imaged using CT-ASLM while **b** and **d** were deparaffinized 50 um sections imaged with LSCM. All scalebars indicate 50 μm.

### Mouse xenograft models of melanoma recapitulate the bleb signaling patterns seen in human cancer patients

To determine whether melanoma xenografts in mice display the same transitions in cell morphology and septin organization at the invasive front of the tumor we used the human MV3 human melanoma cell line used in our previous *in vitro* work^1^. NSG mice received subcutaneous flank injections of MV3 cells, after which tumors were allowed to grow until palpable. Mice were sacrificed, tumors extracted and the tissue cleared for high-resolution imaging by axially scanned lightsheet microscopy (CT-ASLM)^16,17^. This technique allows volumetric imaging at isotropic resolutions of ∼350 nm throughout centimeter-scale tissue samples, making it an ideal tool to study submicron subcellular processes in their larger microenvironmental contexts.

The resulting 3D image volumes recapitulated observations made in patient biopsies. Melanoma cells in the low anchorage conditions characteristic of the invasive margin and stroma possessed blebby amoeboid morphologies with PM-localized septins (Fig 3B, 3D, and Movie 2). In stark contrast, tightly-packed and well-adhered cells within the tumor interior showed non-amoeboid morphotypes that lacked the assembly of septin puncta at the PM (Fig 3C and Movie 2). The visual distinction between bleb signaling active and inactive cells was even more striking than in the biopsy datasets, both because the MV3 cells expressed an endogenous SEPT6-GFP tag that generates clearer separation between diffuse septins in the cytosol and those concentrated in higher-order assemblies at the PM, and because the xenograft tumors possessed far simpler pseudo-spherical geometries, allowing clearer delineation between the invasive margin and tumor interior.

**Figure 3:**
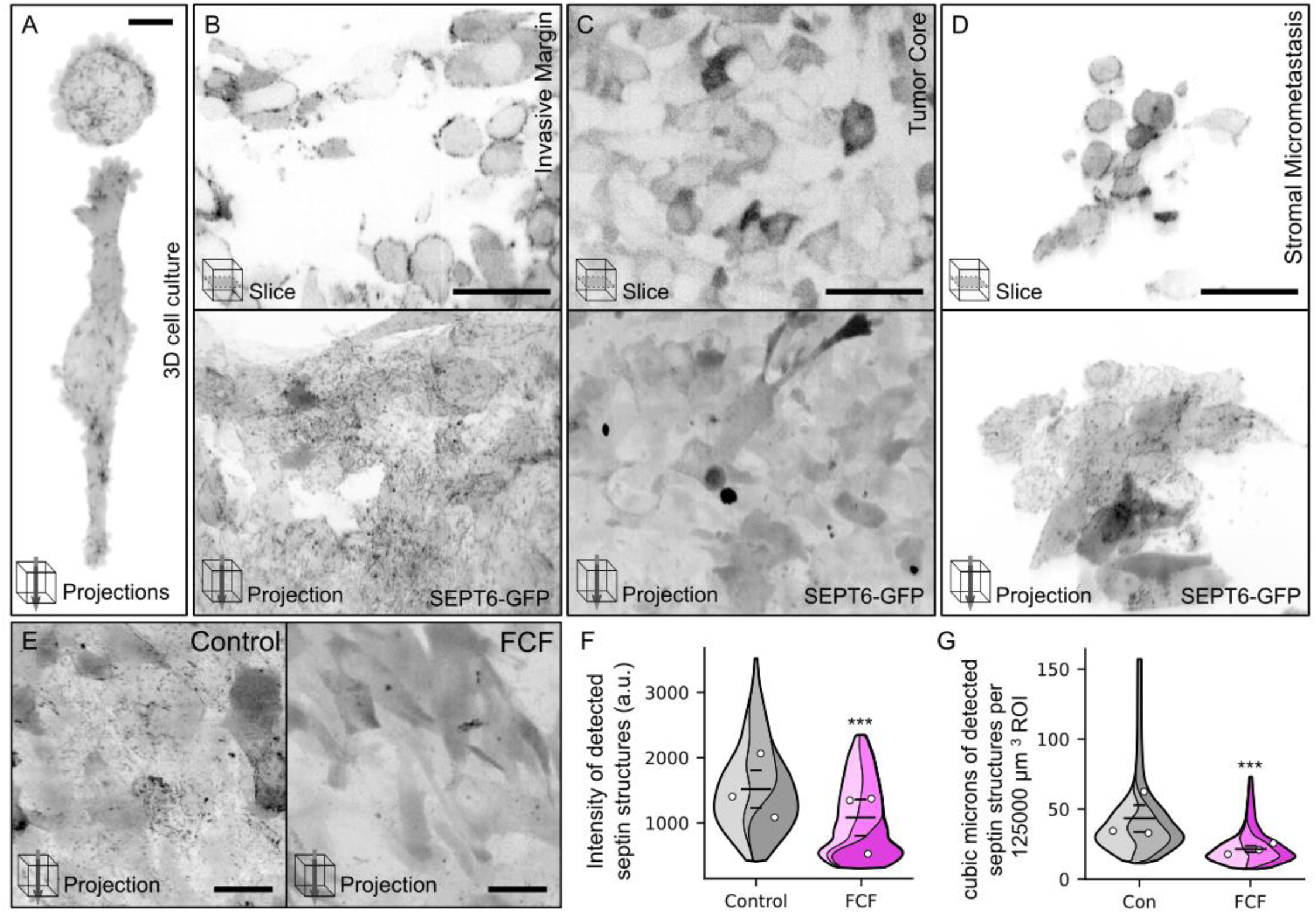
Mouse xenograft models of melanoma recapitulate human *in vivo* results and can be manipulated with a small molecule septin inhibitor. (a) Maximum intensity z-projections of *in vitro* MV3 melanoma cells expressing SEPT6-GFP and imaged while suspended in soft collagen, included for comparison. **(b)** Poorly-adhered MV3 cells showing activation of bleb signaling in the invasive margin surrounding a primary xenografted tumor **(c)** Well-adhered MV3 cells from the interior of the same xenografted tumor shown in **b** displaying no activation of bleb signaling. **(d)** MV3 cells showing activation of bleb signaling in a micrometastasis located in the proximal stroma within 1 mm of a primary xenografted tumor. **(e)** Maximum intensity z-projections of MV3 cells within the invasive margins of xenografted tumors grown in mice either treated with the septin inhibitor FCF (right) or with a vehicle only control (left). **(b-e)** All panels show MV3 cells expressing SEPT6-GFP and grown as xenografted tumors within NSG mice. Tumors were surgically extracted with nearby tissue, chemically cleared, and imaged as intact specimens via CT-ASLM. For **b**, **c**, and **d**, upper panels show single z-slices while lower panels show maximum intensity z-projections. **(f)** Violin SuperPlots of 3 replicate experiments showing the intensity of detected septin structures in cells at the invasive margin of xenografted tumors within treated and untreated mice. **(g)** Violin SuperPlots of the same 3 replicate experiments in **f** showing the total volume of detected septin structures found within individual 125000 um3 subvolumes taken at the invasive margins of xenografted tumors within treated and untreated mice. For **f** and **g**, white circles indicate individual replicate means, long horizontal lines indicate combined means, short horizontal lines indicate combined SEMs, and color-coding is shared between panels. Statistical testing was performed using Welch’s t-test. Scalebars indicate 10 μm for **a**, 50 μm for **b-d**, and 25 μm for **e**.

### Oral administration of the septin inhibitor FCF effectively targets bleb signaling in mouse xenograft tumors

Having established that our experimental system recreated the septin phenotype observed in primary tumor biopsies, we next sought to determine its experimental pliability. To this end, we repeated the xenograft experiments in 6 mice, half of which were treated with the septin inhibitor forchlorfenuron (FCF)^18,19^. FCF has long been used as an effective inhibitor of septin function in both mammalian and fungal cells, and we have previously demonstrated its ability to disrupt PM septin localization and bleb signaling in melanoma cells *in vitro*^1^. While it has only been employed as a septin inhibitor in a very small number of murine studies^19–21^, FCF has been used as an agricultural chemical for over 40 years (where it is often referred to as CPPU or 6-CPPU), resulting in a large number of toxicity studies performed in a variety of mammalian models.^22,23^ All such studies suggest little-to-no acute or chronic toxicity, mutagenicity, or carcinogenicity, especially relative to the effects generally produced by anti-cancer therapeutic compounds.

After tumors became palpable in all 6 mice, we prepared FCF suspensions in carboxymethyl cellulose and treated mice via oral gavage for 3 days. Tumors were then extracted, cleared, and visualized using CT-ASLM as before. We found that FCF treatment had a marked effect on septin localization within cells at the invasive margin, resulting in significantly fewer septin structures at the PM, with those structures that remained displaying significantly reduced fluorescent signal (Fig 3E-G). Thus, we report that FCF is bioavailable via oral gavage and effectively disrupts septin-mediated bleb signaling, just as seen *in vitro*.

### Anti-septin therapy specifically kills melanoma cells within the invasive margin and proximal stroma

We next sought to determine the effect of prolonged septin inhibition on xenografted tumors and cells. Xenografts were grown in 12 mice and, as before, FCF treatments were started in half of the animals after tumors became palpable. These treatments continued in a 5-days-on/2-days-off protocol until the largest tumor reached 1.5 cm in length (21 days), at which point all animals were sacrificed and tumors were recovered. As previous work suggested that FCF is hepatically metabolized^24^, serum was also collected from each animal in order to run metabolic tests of liver function, all of which showed negligible signs of systemic toxicity (Fig S2).

We began by assaying cell death using paraffin embedding and TUNEL staining. While there was no appreciable difference between treatment groups in the overall size of extracted tumors (Fig 4A), tumors from FCF-treated mice produced significantly higher TUNEL signals (Fig 4B). Critically, we consistently observed the TUNEL signal to be concentrated near tumor margins (Fig 4B, inset), consistent with the location of cells containing puncta of assembled septin. Hence, FCF selectively targets the septin scaffolds that are necessary for bleb-induced survival signaling at low-anchorage periphery of tumors. We confirmed this with higher resolution imaging via CT-ASLM, finding little-to-no TUNEL signals at the invasive margins of control tumors, while these same regions showed very high levels of TUNEL staining in FCF-treated mice (Fig 4C and Movie 3). Indeed, this high-resolution visualization shows that TUNEL signals tend to be localized just outside the boundary of the tumor, in agreement with migrating melanoma cells dying shortly after entering the stroma when experiencing conditions that prevent the formation of septin signaling hubs.

**Figure 4:**
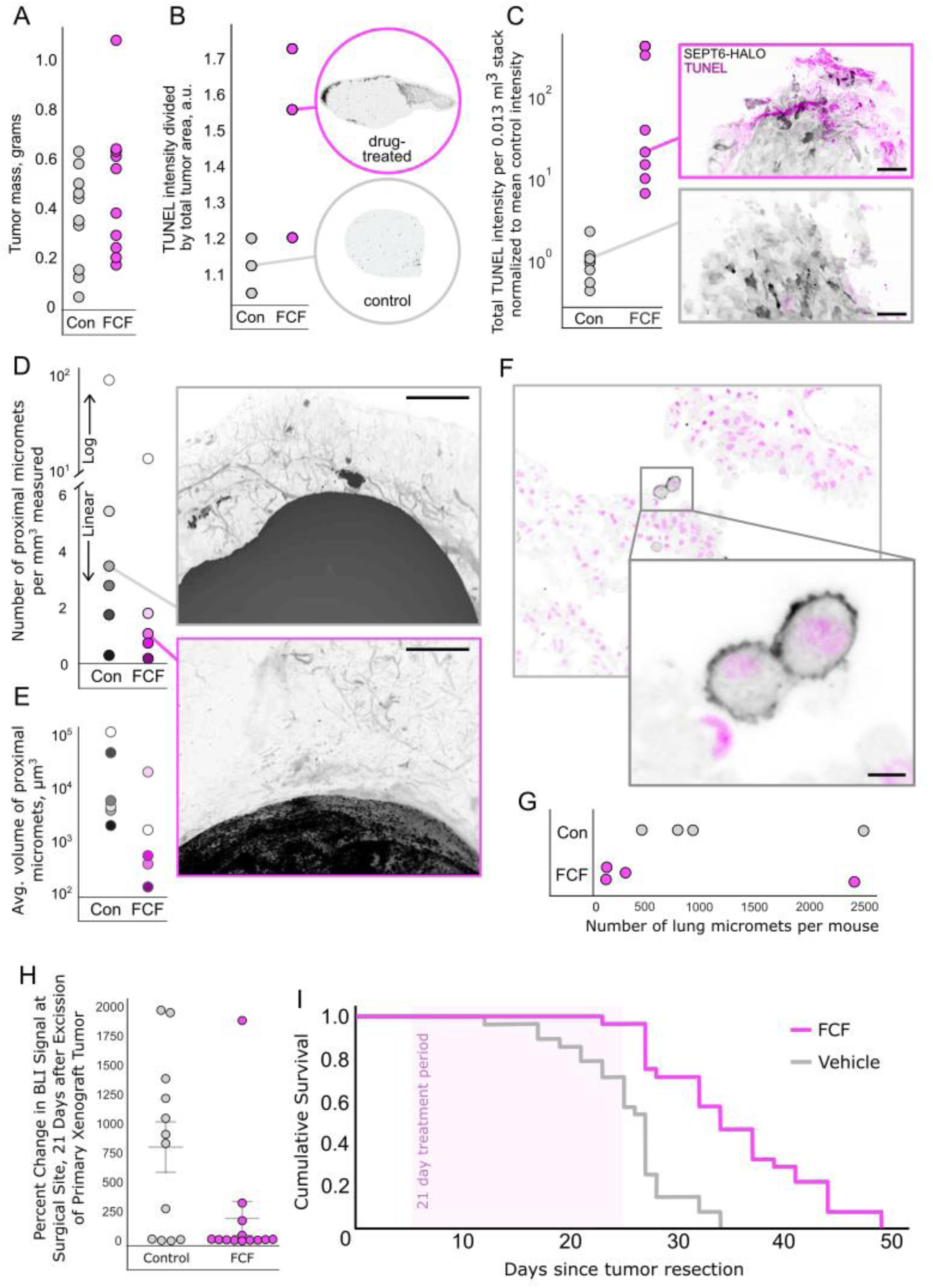
Anti-septin therapy specifically kills isolated and micrometastatic melanoma cells, eliminating proximal and distant micrometastases, preventing tumor recurrence, and prolonging survival. **(a)** The mass of surgically removed tumors from mice that were treated with either FCF or a vehicle-only control for 21 days after tumors became palpable, showing no significant difference. **(b)** Cell death, as measured with TUNEL staining, in surgically removed tumors from 21-day treated and untreated mice. Each data point represents the mean value of 3 medial 20 µm slices from individual tumors. Y-values indicate the total amount of TUNEL intensity (a.u.) measured in the tumor, normalized by area measured. Insets show images from median data points. **(c)** Tumors from 21-day treated and untreated mice were chemically cleared, TUNEL stained, and imaged via CT-ASLM at their invasive margins. Data points represent individual 0.013 mm3 subvolumes collected from 4 different mice. Y-values indicate total TUNEL intensity within each subvolume, normalized to mean control intensity. Insets show maximum intensity z-projections of a single 0.013 mm3 subvolume from median data points with SEPT6-HALO in gray and TUNEL in magenta. **(d)** Tumors and nearby tissue from 21-day treated and untreated mice were chemically cleared, imaged via CT-ASLM, and micrometastases within their proximal stroma were detected. Data points represent the proximal stromata of individual tumors from different mice. Y-values represent the number of detected micrometastases for each data point, normalized by the total volume of stroma measured for that data point. Insets show images from median data points, specifically, centimeter-scale maximum intensity z-projections of tumor-proximal stroma with cancer cells in black (SEPT6-HALO) and stromal tissue in gray (autofluorescence). **(e)** The mean volume of detected micrometastases from the same dataset as **d**, with color coding shared between panels. **(f)** Single z-slice of spontaneous micrometastatic melanoma cells in the lung, arising from subcutaneous flank injections of YUMM 1.7 cells expressing cytoplasmic GFP. Cells were immunostained for GFP, shown in black. Nuclei stained with SYTOX Green, shown in magenta. **(g)** Both lungs from 21-day treated and untreated mice were surgically removed, chemically cleared, and imaged with CT-ASLM. Y-values represent the full count of all micrometastases in both lungs of 8 different mice. **(h)** Tumors were xenografted into 25 mice and allowed to grow until palpable, after which they were surgically excised and mice were treated for 21 days with either FCF or a vehicle-only control. BLI intensity at the surgical site was measured 1 day and 21 days post-surgery, with y-values representing the percent change in these signals for individual mice. Whiskers represent the mean and SEM of each dataset. **(i)** The experiment in **h** was repeated using 56 mice, with a 21-day FCF treatment beginning 5 days after surgery. Plot shows Kaplan-Meier survival curves for each group. Log-rank test, p-value = 0.0000112. Scalebars indicate 70 µm for **c**, 1 mm for **d and e**, and 10 µm

### Anti-septin therapy clears micrometastases from the proximal stroma and sites of distant metastasis

To assess whether this increase in context-specific cell death has an effect on disease progression, we investigated the extent of metastatic spread in the stroma surrounding tumors. In control tumors, the spread almost exclusively took the form of satellite micrometastases – small, mostly punctate, collections of 1-100 cancer cells^25^. While they were relatively rare and sparsely distributed in most animals, a full centimeter-scale volumetric scan revealed dozens of such colonies in the tumor proximal stroma, while FCF-treated animals possessed significantly fewer (Fig 4D-E and Movie 4). Since septin inhibition reduced micrometastatic burden in the proximal stroma, we wondered whether it might produce a similar effect in distant organs. Examination of spontaneous lung micrometastases arising from subcutaneous xenograft tumors confirmed that these isolated tumor cells take on the same blebby amoeboid morphotype observed in the invasive margin, tumor-proximal stroma, and malignant effusions (Figure 4F and Movie 5). To test the effect of septin blockade on these micrometastases, we cleared and imaged the lungs of FCF treated and untreated mice, finding that, just as seen in the stroma, anti-septin therapy also reduced micrometastatic burden in the lungs (Fig 4G and Movie 6). These results support the hypothesis that bleb signaling occurs *in vivo* and possesses the same pro-survival function as it does *in vitro*, and that targeting this pathway meaningfully impairs disease progression by killing metastatic cells.

### Anti-septin therapy effectively prevents tumor recurrence and prolongs survival

All data presented thus far indicate that anti-septin therapy does not target tumors in a general sense but instead specifically kills micrometastatic cells. We sought to design a new series of experiments encompassing the type of clinical scenario in which such a therapy would be valuable. Cancer recurrence is a major cause of mortality in melanoma patients^26,27^ and, as it is driven by residual micrometastases that persist after treatment, its prevention is an excellent candidate for such a therapeutic approach. To this end, we performed experiments on mice in which the primary xenograft was surgically excised after growing to palpability. FCF treatment then began on the day after surgery, after elimination of detectable tumor at the surgical site was confirmed using whole-body bioluminescent imaging (BLI). Half the mice were treated for 3 weeks using the same protocol as before, imaged again and compared to mice that did not receive treatment. These BLI datasets showed that after 21 days, 66% of control mice had detectable local tumor recurrence at the surgical site, compared to only 23% of the mice that received anti-septin therapy (Fig 4H). Moreover, the recurrent tumors in FCF-treated mice produced BLI signals far dimmer than those of recurrent tumors in untreated mice, suggesting a significantly lower tumor burden. We repeated the experiment, treating mice for 3 weeks after surgery with either FCF or a vehicle-only control, this time tracking survival. As expected, mice receiving FCF treatment survived significantly longer, with a mean survival period of 35.2 days vs 25.4 days for untreated mice (Log-rank test, p-value = 1.12 × 10^−5^, Fig 4I).

## DISCUSSION

Our data indicate a role for bleb signaling in cancer disease progression and motivate continued translational work exploring the pathway as a therapeutic vulnerability of metastasizing cancer cells. The data support not only the central hypothesis that bleb signaling is a central pathway for anchorage independent cell survival, but also outlines the clinical blockade of the pathway as a potentially powerful means of adjuvant therapy. Septin inhibition has no appreciable effect on tumor size or growth rates. However, it is very effective at specifically targeting isolated and micrometastatic cancer cells. Thus, such a therapy could be an effective complement to primary treatments such as chemotherapy, radiotherapy, immunotherapy, targeted therapy or surgical intervention, all of which can leave recurrence-enabling persister cells and occult metastasis. This possibility is further bolstered by FCF seeming to possess low toxicity, projecting an increased tolerability of such a treatment in combination with other drugs.

Our previous work showed that the activation of bleb signaling is a highly dynamic and fast-acting process. Melanoma cells begin blebbing immediately upon losing attachment, activating the nascent signaling potential of cytosolic septins by translocating them into hubs at the PM where they replace the signals lost upon the disruption of adhesion-dependent signaling hubs such as focal adhesions, adherens junctions, and desmosomes. The speed with which this process is initiated allows cells to quickly respond to changing microenvironmental conditions and stressors at a rate outpacing transcriptional and translational regulation. It is interesting then that when examined *in vivo*, bleb signaling does not seem to become *inactivated* as readily as it is activated. Indeed, in this study we found evidence of bleb signaling not only at the invasive margin where attachment is initially lost, but also in micrometastatic cells in the stroma, within a variety of malignant effusions, and at sites of distant metastasis. Why then do these cells not form attachments in their new microenvironments and accordingly inactivate bleb signaling? It is possible that in all these locales they are unable to do so (perhaps due to a lack of molecular substrates suitable for the adhesion factors they express) and thus are experiencing consistent microenvironmental cues which continually reinforce the amoeboid bleb signaling phenotype. It is also possible that once this phenotype is adopted cells become broadly reliant upon it, similar to oncogene addiction. Regardless of the underlying mechanism, this durable activation of bleb signaling seems to result in the sensitization of disseminated melanoma cells to septin inhibition across a broad diversity of microenvironments. It would be worthwhile to study whether this behavior is related to metastatic organotropism, if a shift away from bleb signaling coincides with the reactivation of these dormant cancer cells, and whether known reactivating signals such as inflammation can trigger this phenotypic shift.

Though we made use of FCF in this study, the best pharmacological means of inhibiting bleb signaling in a preclinical experimental context remains an open question. However, the specifics of the pathway suggest septins are the ideal target. This owes to their seemingly unique ability to respond to the upstream activating signal of bleb-generated micron-scale membrane curvature, making them the pathway’s sole known molecular bottleneck^1^. FCF has been used as a small molecule septin inhibitor in research for decades^18,19^, and the results shared here suggest it is both effective and safe (a characteristic that is backed by 20+ years of FDA and EFSA studies demonstrating low toxicities in animal trials^22,23^). However, any eventual clinical usage would be biologically complicated by the suggestion of potential off-target mitochondrial effects^28^ (now somewhat mitigated by an increased understanding of septins’ role in mitochondrial function^29–31^), and financially complicated by the fact that it has aged out of its initial patents, making it difficult to fund clinical trials exploring its efficacy and safety in cancer patients. At least two different groups have developed FCF derivatives that improve binding affinity to produce more effective septin inhibition^32,33^, but unfortunately this results in an accompanying increase in toxicity, especially at organismal scales. UR214-9 is arguably the most promising of these derivatives, though recent studies indicate it is still unacceptably toxic relative to FCF^30^. These results raise the possibility that FCF’s specific binding profile might represent an optimal balance of efficacy and safety, with action sufficient to disrupt septin hub formation, yet still below the activity threshold necessary for broad toxicity. Other potential avenues exist, such as new septin-targeting compounds developed by Remynd. Chemically unrelated to FCF, these compounds act as a “molecular glue” increasing septin polymerization with the goal of restoring calcium signaling in dysfunctional neurons^34–36^. It will be interesting to test whether stabilization of septin assembly has the same disruptive effect on bleb signaling as blocking assembly, especially should these or similar compounds prove safe for human use.

The experiments described here have an explicit focus on bleb signaling in melanoma, a choice made to expand on our previous *in vitro* work relying on melanoma cell lines as model systems, seeking compatible results by generating analogous clinical and xenograft *in vivo* datasets. Despite the limitation to one cancer type, our data shows hints of a potential universality of bleb signaling in cancer. This is most apparently seen in Figure 2C, which strongly indicates the occurrence of bleb signaling in Merkel cell carcinoma (MCC). This is significant because, though it is often categorized as a cancer of the skin, MCC has a distinct origin and etiology from melanoma^37^. This prompts the question of what other non-melanoma cancer types might employ bleb signaling and raises the possibility that anti-septin therapeutics targeting the pathway might possess broad efficacy. Such breadth is further supported by molecular evidence that septin hubs are scaffolding HER2 in the mucosal melanoma cytology dataset (a phenomenon previously shown to be central to the survival of gastric cancer cells *in vitro*^13^) (Figures 1b, 1d, S1 and Movie 1). Our previous *in vitro* dataset in MV3 and other melanoma cell lines displayed scaffolding of NRAS rather than HER2 leading to upregulation of MAPK and PI3K^1^. This means that not only do septin hubs appear capable of scaffolding different oncogenic pathways in different cancers, but also that the pathways in question are the specific drivers of those cancers. The apparent promiscuity is not especially surprising, as septins have been shown to scaffold dozens of signaling pathways across multiple kingdoms of life^38–40^, but the implication of the centrality of septin hubs and bleb signaling to the pathophysiology of different cancer types increases the potential for clinical versatility. Though these conclusions are clearly speculative due to the limited amount of data currently available, we feel the inherent impact of the suggestion makes them deserving of both discussion and future experimental work.

Altogether, these findings highlight bleb signaling as a mechanistically distinct and therapeutically actionable feature of metastatic cancer, with septin hubs emerging as a promising molecular target whose inhibition may complement existing treatment modalities and broaden clinical impact across diverse cancer types. The hypothesis-driven discovery of this non-canonical oncogenic behavior has yielded the identification of a morphologically-regulated cellular process which, though it is relatively rare and environment-specific, nonetheless seems to be instrumental in the progression of melanoma. It is important to consider that bleb signaling’s activation, in which cancer cells selectively and transiently leverage a rapid reorganization of their biochemical regulation, is independent of the kinds of genetic and transcriptional adaptations that would make it detectable by genomic and transcriptomic datasets. This underlines the value of understanding the complexities of proteomic signaling events at the plasma membrane and how those events can be regulated by cell morphological programs and similarly overlooked cellular states. Moreover, it demonstrates the continued importance of hypothesis-driven cell biology in the discovery of novel cancer vulnerabilities, even in an era when large-scale acquisition of multi-factorial datasets and unbiased exploration are the dominant approach.

## METHODS

### Cytology Clinical Samples

Cytology specimens were collected from the Brigham and Women’s Hospital and collected under Institutional Review Board approval (FWA00007071, Protocol IRB18-1363) under a waiver of consent. The mucosal melanoma sample was obtained via peritoneal fluid collection from a 51 y.o. male patient diagnosed with metastatic anal mucosal melanoma. The cutaneous melanoma sample was obtained via fine needle aspiration of a salivary gland lesion from a 62 y.o. male patient diagnosed with Stage IIC BRAF(V600E) melanoma originating from the skin of the right forearm. The vulvar melanoma sample was obtained via pleural fluid collection from a 65 y.o. female patient diagnosed with metastatic malignant melanoma originating form the skin of the left vulva. All samples were identified by a board-certified pathologist through immunohistochemical staining for SOX10 and MART1 and/or melanoma diagnostic criteria based on H&E staining.

For the immunofluorescent study of the mucosal melanoma sample, a 20 µm section was cut, dewaxed, and stained using the protocol outlined in Yapp, et al., 2025^41^. FFPE specimens were dewaxed on a Leica Bond Rx. Samples were then incubated for 8 hours at room temperature with a rabbit primary antibody against SEPT2 (abcam, ab187654) followed by a goat anti-rabbit secondary antibody conjugated to AlexaFluor555 (Invitrogen, A21430) for the same span. In between these incubations the sample received 3 PBS washes for 1 hour each. The sample was washed again and incubated with a primary antibody against HER2 conjugated to AlexaFluor647 (abcam, ab225510) for 8 hours at room temperature. It was washed again, stained with Hoechst 33342 (ThermoFisher) in Superblock buffer (ThermoFisher) at 4 degrees overnight followed by a final wash. Samples were then mounted with 70% glycerol under a #1.5H coverslip (Thor labs) and imaged with confocal microscopy as described below.

### Primary Tumor Clinical Samples

The nodular acral melanoma and Merkel cell carcinoma specimens were procured from the Cooperative Human Tissue Network (CHTN, Eastern Division, University of Pennsylvania, Philadelphia, PA, USA) in accordance with institutional and federal ethical guidelines. All tissues were de-identified and supplied under a Human Subjects Agreement prohibiting re-identification or unauthorized use. Based on institutional review, this study was determined not to constitute human-subjects research and thus did not require IRB oversight. The Merkel cell carcinoma specimen originated from a wide excision of the left thigh, measuring 6.5 cm in maximum dimension, with multifocal lymphovascular invasion and negative deep margins. The nodular melanoma specimen was excised from the right lateral breast and exhibited a Breslow thickness of 45 mm, Clark level V, and extensive nodal metastasis involving 13 of 24 lymph nodes. Both specimens were fixed in 4% PFA after excision.

Tissues were shipped on dry ice and stored at 4°C upon arrival until processing. All specimen handling adhered to OSHA standards for bloodborne pathogens and institutional biosafety protocols. Samples were then incubated in primary antibodies: SEPT2 diluted (1:100) in staining buffer for 72 hours at room temperature, shielded from light. After primary labeling, tissues were washed in Wash Buffer (PBS + 0.5% NP40, 10% DMSO) with buffer changes every 2 hours for 6 hours. The samples were post-fixed overnight in 4% PFA. Following post-fixation, tissues were washed with PBS and incubated in secondary antibody (Donkey anti-rabbit AF 488, 1:2000) in staining buffer for 72 hours at room temperature, protected from light. After secondary labeling, tissues were washed with Wash Buffer every 2 hours for 6 hours and left in Wash Buffer overnight. The next day, Wash Buffer was refreshed.

The superficial cutaneous melanoma and mucosal melanoma samples were collected from the Brigham and Women’s Hospital and collected under Institutional Review Board approval (FWA00007071, Protocol IRB18-1363) under a waiver of consent. The superficial cutaneous primary melanoma was from a 31 y.o. female patient diagnosed with FFPE Stage III melanoma. The primary mucosal melanoma sample was obtained from the same 51 y.o. male patient described in the cytology methods above. Both samples were identified by a board-certified pathologist through immunohistochemical staining for SOX10 and MART1 and/or melanoma diagnostic criteria based on H&E staining.

35-micron thick samples were cut, dewaxed, and stained using the protocol outlined in Yapp, et al., 2025^41^. FFPE specimens were dewaxed on a Leica Bond Rx and bleached to remove autofluorescence using the standard CyCIF protocol^42^. For bleaching, samples were submerged in 5ml H2O2, 800ul NaOH, 25ml PBS between LED light panels for 1 hour. Then, samples were incubated with primary conjugated antibodies and Hoechst 33342 (ThermoFisher) in Superblock buffer (ThermoFisher) at 4 degrees overnight followed by washes in PBS for 3 times for 1 hour at room temperature. Samples were then mounted with 70% glycerol under a #1.5H coverslip (Thor labs).

### Mouse Xenograft Studies

#### Septin Localization Studies

To analyze the regional context of bleb signaling activation in tumors and proximal tissue (Fig 3B-G), MV3 human melanoma cells carrying SEPT6-eGFP^1^ were used to generate xenograft tumors. 2 million melanoma cells in 100 mL PBS were injected into the subcutaneous space of the right flank of NOD-SCID-Il2rg−/− (NSG) mice. All mice were engrafted and formed tumors. Once they were approximately 1-1.5 cm in diameter, tumors were surgically resected along with nearby tissue, after which mice were euthanized. Pilot studies showed GFP signal to be maintained after clearing with CUBIC, though unstable. For this reason, all datasets were collected within 48 hours of completion of the CUBIC protocol (see below).

#### FCF Treatment

Anti-septin treatment was performed via oral gavage of 10 mg/ml FCF suspended in 0.5% carboxymethyl cellulose (CMC). Study animals received 10 µl of suspension per gram of body mass in a 5-days-on/2-days-off protocol until tumors reached 1.5 cm in length along their longest axis, after which tumors were harvested. Control animals received the same volume of 0.5% CMC alone.

#### Disease Progression Studies

For initial disease progression studies (Fig 4A-F), the above mouse protocols were followed, with the exception of the cell line used. MV3 cells carrying SEPT6-HALO were used instead of SEPT6-eGFP due to the aforementioned instability of GFP signal post-clearing. After tumor excision, the samples were weighed and then fixed with gentle agitation for 20 minutes with 4% PFA in 1x PBS at 4°C. The fixative solution was removed with two washes of 5 minutes with 1x PBS. Then the samples were incubated with gentle agitation in HALO TMR ligand (1uM) for 2 hours at room temperature^43^. Tumors were washed twice for 5 minutes with 1X PBS to remove the ligand. To complete the tumors fixation after the Halo TMR ligand staining, the samples were incubated again in 4% PFA in PBS at 4°C overnight.

Blood was also harvested from all animals via cardiac puncture. Clotted blood was spun at 15,000 RPM for 10 mins at 4 °C, and the supernatant was collected as serum. ALB, ALKP, ALT, AST, TBIL, and LDH levels were measured with the VITROS 350 clinical analyzer (microchemical slides technology, Ortho Clinical Diagnostics, Raritan, NJ).

#### Histological Cell Death Assay

Initial TUNEL analysis (Fig 4b) was performed by UT Southwestern’s Histo Pathology Core according to standard procedures^44,45^. Samples were dehydrated through 7 increasing grades of ethanol, cleared through 3 exchanges of xylene and infiltrated with 3 exchanges of paraffin wax (Paraplast Plus, Leica Microsystems, Deer Park, IL, USA) with vacuum-assist on a Thermo-Shandon Excelsior Tissue Processor (Kalamazoo, MI, USA). Embeds to obtain tumor anatomy inclusive of capsule contiguous to core were prepared on a Sakura Finetek Tec 6 Embedding Center (Torrance, CA, USA). Serial paraffin sections were prepared and checked by dark-field microscopy to ensure quality without rarefaction. Positive nuclei of apoptotic and necrotic cells were labeled with fluorescein according to methods of first report^44^ and literature supplied with the DeadEnd Fluorometric TUNEL System. Nuclei were counterstained with propidium iodide and coverslipping performed using Vectashield (Vector Laboratories, Cat# H1000).

#### Cleared Tissue Cell Death Assay

Click-iT TUNEL Alexa Fluor Imaging Assay from Invitrogen (C10247) was used to assay cell death in cleared tumor samples. After fixation the samples were washed for 6 hours with PBS 0.02% sodium azide refreshed every 2 hours. Then tumors were cut in ∼2mm slices using a tissue matrix slicer (Prod No 15020, 0.5mm slices fromTED PELLA,INC). Tumor slices were permeabilized with 0.25% Triton-X-100 buffer with gentle agitation overnight at 4°C. After tumors permeabilization the TdT reaction was performed. Samples were incubated in the TdT reaction buffer (component A) for 30 minutes at room temperature with gentle agitation and then removed. The TdT reaction cocktail was prepared according to Invitrogen *User Guide* and the samples were incubated in the TdT reaction cocktail overnight at RT with gentle agitation. The next day the samples were washed twice with 3% BSA in PBS for 5 minutes. Then, the Click-iT reaction was performed. Tumors were incubated in the Click-iT reaction cocktail (prepared as the User Guide) for 1 hour with gentle agitation at RT. The Click-iT reaction cocktail was removed with two washes with 3% BSA in PBS for 5 minutes each.

#### Tumor Recurrence Study

MV3 melanoma cells used in this study expressed luciferase so that disease burden could be determined by bioluminescence imaging. Tumors were formed as above. When the tumors reached approximately 1-1.5 cm in diameter, they were surgically resected, post-surgical bioluminescence imaging was performed, and mice began receiving FCF as above. Three weeks after the treatment, bioluminescence imaging was performed again to assess the effects of the drug on local recurrence at the surgical site.

Bioluminescence imaging was performed as previously described (Piskounova et al., Nature 2015). Briefly, mice were injected intraperitoneally with 100 μl of PBS containing D-luciferin monopotassium salt (40 μg ml−1) (Goldbio Cat# LUCK-3G) 5 min before imaging, followed by general anesthesia 2 min before imaging.

#### Post-Resection Survival Study

To test the effects of the septin inhibitor on survival after primary tumor resection, 100 cells in 50 mL of 25% growth factor-reduced Matrigel (Corning Cat #354230) were injected into the right flank of NSG mice. This alternative method was used to minimize the possibility of multiple primary xenografts being engrafted due to larger injection volumes or higher numbers of injected cells. 57 out of 60 mice engrafted and formed primary tumors. When the tumors reached approximately 1-1.5cm in diameter, we surgically resected the tumors and started treating the mice 5 days after surgery with FCF as above. Survivorship was measured daily and Kaplan-Meier survival analysis was performed on the resulting data.

### CUBIC tissue clearing

Following xenograft tumor harvesting, samples were immediately fixed with 4% paraformaldehyde (PFA) at 4°C overnight. Fixed tissues were then washed with PBS containing 0.02% sodium azide at room temperature with gentle agitation for 6 hours, the PBS was refreshed every 2 hours. Samples were left overnight in the PBS with gentle shaking. The next day the samples were pre-treated with a 1:1 mixture of water and CUBIC-L (50% CUBIC-L) at 37°C with gentle shaking for 6-24 hours approximately. The next day the tumors were delipidated with 100% CUBIC-L at 37°C with gentle shaking. The 100% CUBIC-L was refreshed after overnight incubation and then every 2 days until no supernatant color change was visible. After delipidation, samples were washed three times in PBS for a total of 6 hours, replacing the PBS every 2 hours. To initiate refractive index (RI) matching, samples were incubated at room temperature in 50% CUBIC-R+ (1:1 dilution with water) for 24 hours with gentle shaking, followed by incubation in 100% CUBIC-R+ at room temperature until imaging.

### BABB tissue clearing

Mouse lungs and tumors, and human primary tumor samples were cleared using a modified BABB protocol. All steps were performed with gentle rotation, and samples were protected from light. Freshly collected tissues were fixed in 4% PFA at 4°C for less than 24 hours. After fixation, tissues were washed in PBS with 0.02% sodium azide three times for 2 hours each, followed by overnight incubation at 4°C. Tissues were then cut into ∼1-2 mm slices using a matrix and placed in individual 1.5 mL tubes. Pre-treatment involved incubating the samples in 25% QUADROL at 37°C with shaking, and non-perfused tissues were refreshed until the supernatant was clear. Perfused samples were incubated overnight in 25% Quadrol. Following Quadrol treatment, tissues were washed with PBS and incubated in blocking buffer (PBS + 0.5% NP40, 10% DMSO, 5% serum, 0.5% Triton X-100) for 1 day at room temperature. Samples were then incubated in primary antibody diluted in staining buffer for 72 hours at room temperature, shielded from light. After primary labeling, tissues were washed in Wash Buffer (PBS + 0.5% NP40, 10% DMSO) with buffer changes every 2 hours for 6 hours. The samples were post-fixed overnight in 4% PFA. Following post-fixation, tissues were washed with PBS and incubated in secondary antibody in staining buffer for 72 hours at room temperature, protected from light. After secondary labeling, tissues were washed with Wash Buffer every 2 hours for 6 hours and left in Wash Buffer overnight. The next day, Wash Buffer was refreshed. For nuclear staining, tissues were incubated in a dye such as SYTOX Green 488 (1 µM) for 48 hours at room temperature. After staining, tissues were washed twice with PBS for 15 minutes and dehydrated in a methanol gradient (25%, 50%, 75%) for at least 1 hour per step. The samples were then placed in 100% methanol for 45 minutes, refreshed, and rotated for an additional 45 minutes. After methanol dehydration, the tissues were delipidated with two 30–45-minute washes in dichloromethane (DCM). Samples were then incubated in fresh BABB (1:2 benzyl alcohol: benzyl benzoate) for tissue clearing, with three buffer changes. The cleared samples were incubated overnight in fresh BABB at room temperature, protected from light. All BABB solutions were prepared by placing 45mL of BABB in a 50mL conical with 5g of activated aluminum oxide (to remove peroxides), gently agitating the solution for at least 1 hour at room temperature and centrifuging the tube at 3000rpm for 10 mins to pellet the aluminum oxide.

### Confocal Microscopy

Imaging was performed on a LSM980 (Carl Zeiss) laser scanning confocal microscope fitted with a 10x/0.45 NA air and 63x/1.4 NA oil immersion objective lens running ZEN 3.7. An overview scan of both samples was first obtained at 10x magnification. From these preview images, we identified ROIs for further cycling at 63x with z-stacks (sampled at 132nm in x,y; 250nm in z for the cytology specimen and 90nm in x,y; 190nm in z for the primary melanoma). Pinhole was set to approximately 1 Airy unit. To increase throughput, bi-directional scanning without averaging was used. In addition, channels were separated into two tracks to spectrally separate channels (Track 1 – Hoechst, Alexafluor 555, Alexafluor 750; Track 2 – Alexafluor 488, Alexafluor 647) while allowing for some simultaneous emission collection. Individual single-tiled ROIs and multi-tiled regions were used for the cytology and primary melanoma respectively. Focus support points were used to maintain focus across the sample. Datasets were denoised using LSM Plus and underwent stitching in ZEN 3.7 based on the Hoechst channel as reference channel.

### Lightsheet Microscopy

Volumetric data were acquired on two cleared-tissue light-sheet platforms. For sub-cellular resolution we used a high-NA Axially Swept Light-Sheet Microscope (ASLM) adapted from Chakraborty et al. Three continuous-wave Coherent OBIS lasers (488, 561 and 647 nm) were merged, spatially filtered and five-fold expanded to a 16.5 mm beam. A cylindrical lens created a line focus that was swept laterally by a 4 kHz resonant galvanometer (CRS 4KHz, Novanta) while a voice-coil-mounted mirror (Equipment Solutions, Inc.) synchronously refocused the sheet along its propagation axis, matching the rolling-shutter of a liquid-cooled sCMOS camera (ORCA Fusion BT, Hamamatsu Photonics). Excitation and detection employed identical 0.7-NA long-working-distance objectives (54-12-8, Applied Scientific Immersion) immersed in clearing medium; emission was passed through interchangeable green, red or far-red filters before detection. Z-stacks were collected in sample-scanning mode with a 200 µm piezo (MODEL INFORMATION, PiezoJena) coupled to a motorized XYZ stage (MP-285, Sutter Instrument Company).

Whole-organ imaging was performed on a macroscale ASLM module (Lin et al.) derived from mesoSPIM. Light from the laser launch was collimated (75 mm FL), passed through an electro-tunable lens (EL-15-40-TC-VIS-5D-C, Optotune) to impose dynamic defocus, relayed to a 10 mm galvanometer at the pupil of a modified 50 mm f/1.4 DSLR lens, and digitally scanned to form an axially swept sheet. Fluorescence was collected with an Olympus MVX10 macroscope fitted with a 0.25-NA, motorized 0.63–6.3× zoom, filtered (green, red, far-red) and imaged with a CMOS camera (ORCA Fusion BT, Hamamatsu Photonics). Illumination and detection beams entered a shared sample immersion chamber through 50 mm fused-silica windows; the detection arm was mounted on a precision long-travel stage for focus adjustment. All hardware timing, scanning and acquisition were coordinated by the open source navigate control software.

### Image Analysis

#### Septin Hub Analysis

The invasive margins of tumors from both treated and untreated mice were imaged using CT-ASLM, with the aim to capture regions possessing septin-activated cells. After being blinded to which treatment group the datasets were from, cubical ROIs of 125,000 µm^3^ were constructed around regions containing the most septin active cells in each image stack using ImageJ^45,46^ (totaling 65 FCF-treated ROIs across 3 mice and 77 control ROIs across three mice). To detect septin structures the stacks were background subtracted using rolling ball with a 5 pixel radius, binarized using the maximum entropy threshold algorithm (using the whole stack histogram), and despeckled using a 3×3 pixel median filter. ROIs of individual septin hubs were generated from the resulting binary stack using the ImageJ Analyze Particles function. The total volume of detected hubs was measured in each 125,000 µm^3^ stack. Because many of the septin activated cells were on the edge of the tumor (and thus were found in ROIs containing many non-cellular voxels), this volume of detected hubs was normalized by the ratio of cellular voxels to total voxels within the stack. This cellular volume was calculated by running a rolling ball median filter with a 10 pixel radius on the stack, binarizing using the minimum error threshold algorithm (using the whole stack histogram), and measuring the volume of detected voxels. The normalized volumes of detected hubs within each 125,000 µm^3^ ROI were then visualized using violin superplots^47^ to communicate heterogeneity across mouse replicates. The mean intensity of detected hubs within each ROI was also measured after subtracting background signal from the raw stacks as above.

#### TUNEL Analysis

For quantification of TUNEL signal in histopathological samples, datasets were blinded and ROIs were drawn at the edges of tumors to capture total tumor area. Background signal was subtracted using a rolling ball algorithm with a 25 pixel radius and total TUNEL signal was measured within each whole-tumor ROI. Total intensity was normalized by total tumor area for each sample. For quantification of TUNEL signal in cleared tissue, CT-ASLM stacks of equivalent size were taken at the invasive margins of tumors from FCF treated and control mice. These regions were found using SEPT6-HALO signal alone to prevent experimenter bias. Background subtraction of TUNEL signal was performed with a 25 pixel radius rolling ball algorithm and total TUNEL signal was measured in each stack. All operations performed with ImageJ.

#### Stromal Micrometastasis Analysis

Image stacks of tumors and surrounding tissue were background subtracted using a rolling ball algorithm with a 25 pixel radius. The stromal tissue around imaged tumors was divided into uniformly sized cubical ROIs of approximately 0.068 mm^3^ by manual selection using ImageJ. These ROIs were manually thresholded to select metastases while excluding autofluorescent tissue. Individual micrometastases were detected from these binarized stacks using the ImageJ plugin “3D Objects Counter”. The total number of detected micrometastases were normalized by the total volume captured in ROIs and reported in Figure 4D, while the average volume micromet volume for each sample was reported in Figure 4E.

#### Lung Micrometastasis Counting

Maximum intensity projections of both lungs from all 8 study mice were generated and micrometastases were manually counted by a blinded analyst.

## ACKNOWLEDGEMENTS

The authors would like to thank Anushka Deshmukh for creating the graphical abstract, Zach Marin for assistance with microscopy, Sean Morrison for assistance with mouse studies, Noelle Williams for assistance formulating FCF suspensions, and John Shelton for assistance with pathological analysis of mouse samples.

## SUPPLEMENTAL FIGURES

Movie 1: HER2 scaffolding.mp4

Movie 2: septin localization in xenograft tumors.mp4

Movie 3: Inv Margin TUNEL.mp4

Movie 4: proximal stroma mesospim.mp4

Movie 5: Lung Micrometastases, Cellular Scale.mp4

Movie 6: Effect of FCF Treatment on Lung Mets.mp4

**Supplemental Figure 1:**
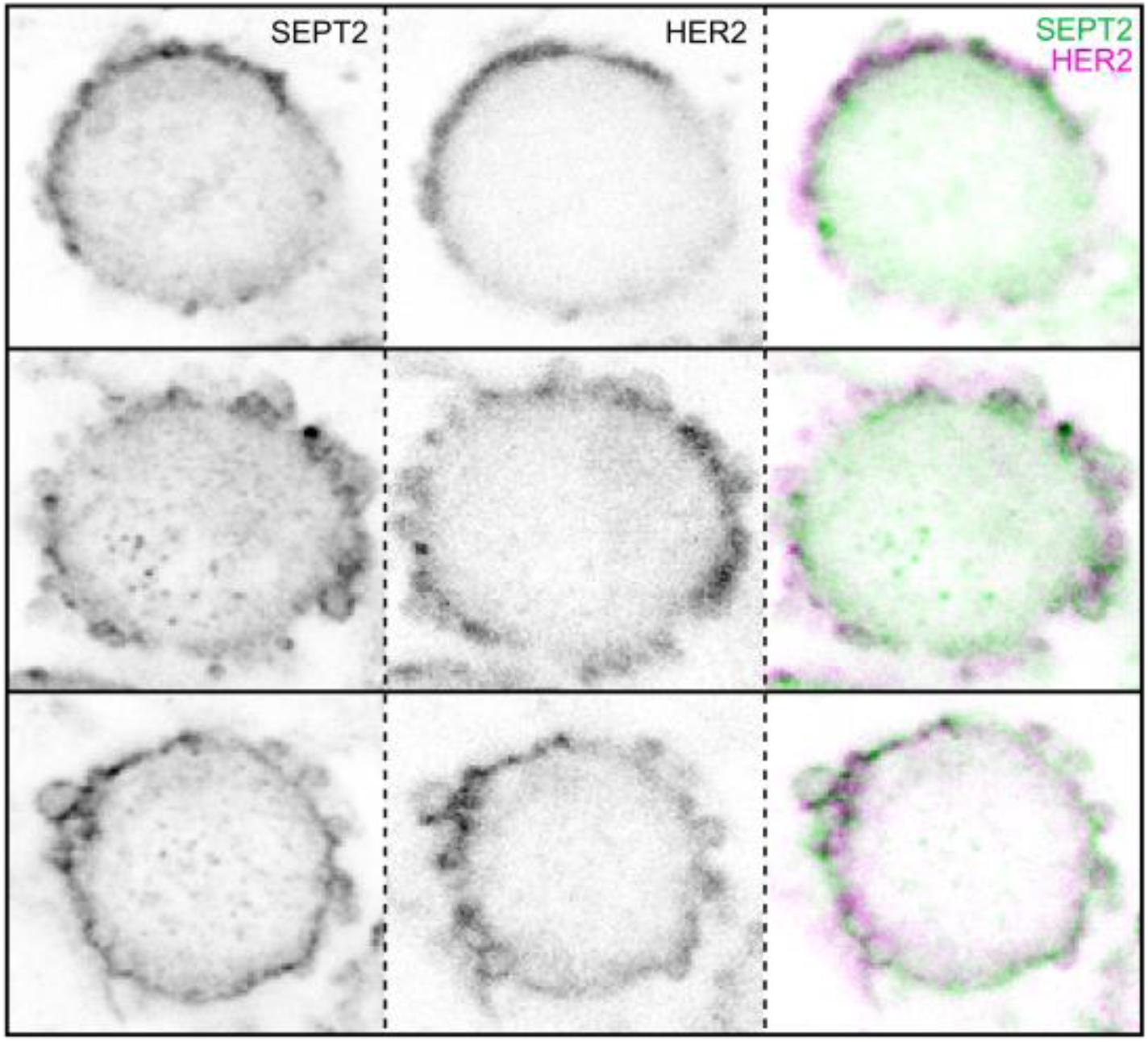
Colocalization data suggest scaffolding of HER2 by septin signaling hubs in metastatic melanoma cells. The same cells shown in Figure 1D, with addition of HER2 immunofluorescence localization data.

**Supplemental Figure 2:**
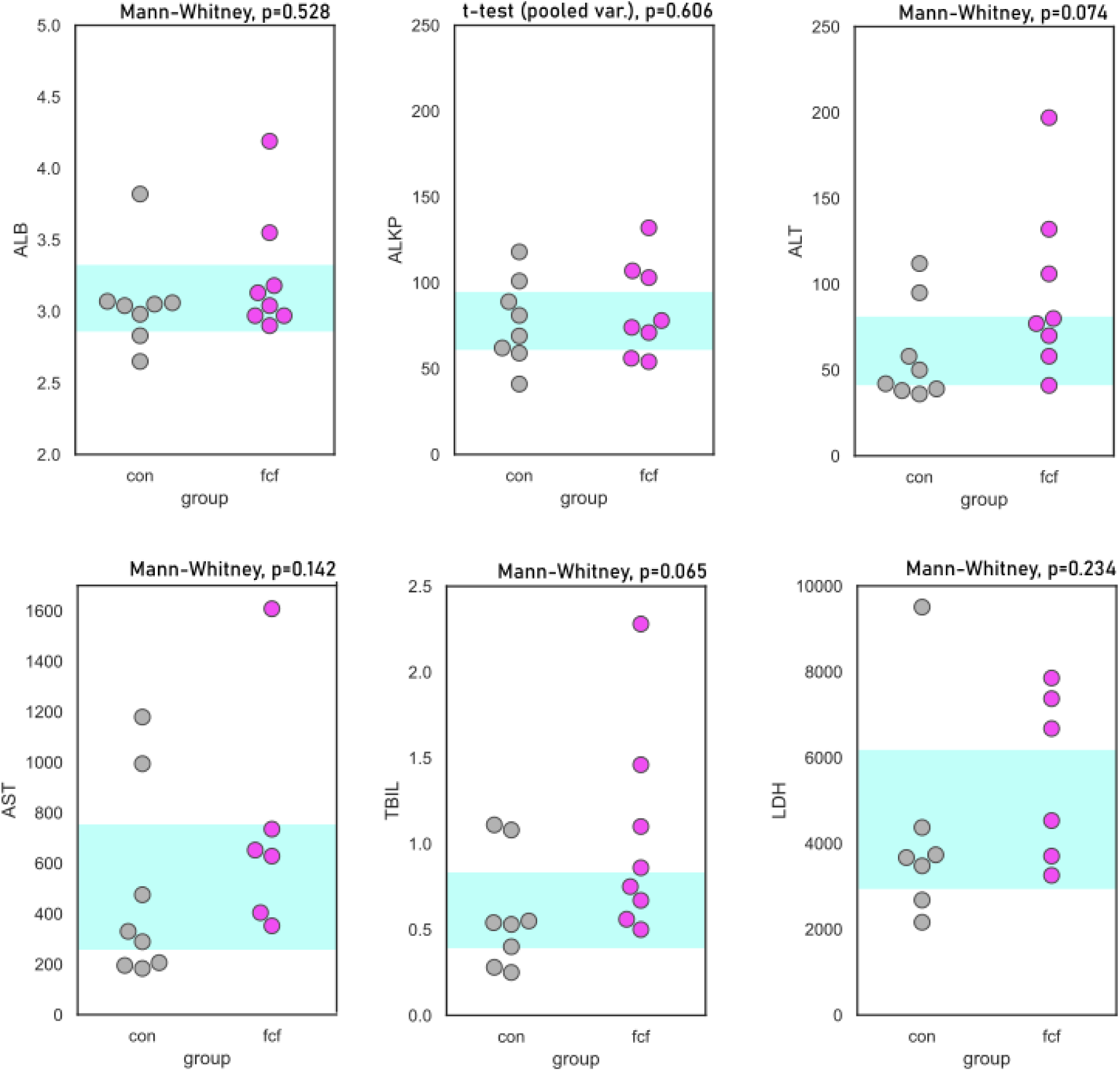
Liver function tests suggest negligible hepatotoxicity. Blood was collected from mice receiving either FCF treatment or vehicle only control for 3 weeks in a 5-days-on, 2-days-off regimen and metabolic tests of liver function were performed. Significance was tested either by Mann-Whitney or t-test, depending on whether F-tests indicated equal or unequal variance (p-values shown above). 95% CI of control group shown in blue band.

